# Antimalarial pantothenamide metabolites target acetyl-CoA synthesis in *Plasmodium falciparum*

**DOI:** 10.1101/256669

**Authors:** Joost Schalkwijk, Erik L. Allman, Patrick A.M. Jansen, Laura E. de Vries, Suzanne Jackowski, Peter N. M. Botman, Christien A. Beuckens-Schortinghuis, Karin M.J. Koolen, J. M. Bolscher, Martijn W. Vos, Karen Miller, Stacy A. Reeves, Helmi Pett, Graham Trevitt, Sergio Wittlin, Christian Scheurer, Sibylle Sax, Christoph Fischli, Gabrielle Josling, Taco W.A. Kooij, Roger Bonnert, Brice Campo, Richard H. Blaauw, Floris P.J.T. Rutjes, Robert W. Sauerwein, Manuel Llinás, Pedro H.H. Hermkens, Koen J. Dechering

**Affiliations:** Radboud University Medical Center, Nijmegen, The Netherlands; Department of Biochemistry & Molecular Biology and Huck Center for Malaria Research, The Pennsylvania State University, University Park, PA 16802 United States of America; St. Jude Children’s Research Hospital, Memphis (TN), United States of America; Chiralix, Nijmegen, The Netherlands; TropIQ Health Sciences, Nijmegen, The Netherlands; XenoGesis Ltd, Nottingham, United Kingdom; Swiss Tropical and Public Health Institute, Basel, Switzerland; University of Basel, Basel, Switzerland; Medicines for Malaria Venture, Geneva, Switzerland; Radboud University, Nijmegen, The Netherlands; Department of Chemistry, The Pennsylvania State University, University Park, PA 16802 United States of America; Hermkens Pharma Consultancy, Oss, The Netherlands

## Abstract

Malaria eradication is critically dependent on novel drugs that target resistant *Plasmodium* parasites and block transmission of the disease. Here we report the discovery of potent pantothenamide bioisosteres that are active against blood-stage *P. falciparum* and also block onward mosquito transmission. These compounds are resistant to degradation by serum pantetheinases, show favorable pharmacokinetic properties and clear parasites in a humanized rodent infection model. Metabolomics revealed that CoA biosynthetic enzymes convert pantothenamides into drug-conjugates that interfere with parasite acetyl-CoA anabolism. *In vitro* generated resistant parasites showed mutations in acetyl-CoA synthetase and acyl-CoA synthetase 11, confirming the key roles of these enzymes in the sensitivity to pantothenamides. These new pantothenamides provide a promising class of antimalarial drugs with a unique mode of action.

**One sentence summary:** Pantothenamides form antimetabolites that interfere with acetyl-CoA metabolism in the human malaria parasite *Plasmodium falciparum*

## Introduction

Malaria is a major global infectious disease that causes approximately 420,000 deaths per year, predominantly among children and pregnant women in sub-Saharan Africa (*1*). It is a vector-borne disease, and transmission depends on the transfer of sexually differentiated gametocytes from human peripheral blood to a mosquito, while the clinical disease is caused by cyclic asexual parasite replication in red blood cells. Eradication of malaria is threatened by the ever-present rise of clinical resistance to current drugs, including artemisinin derivatives, which calls for the urgent development of new antimalarial drugs (*2*). Ideally, new antimalarials should have a novel mechanism of action, no cross-resistance to existing drugs, and demonstrate activity against both asexual stages and sexual blood stages to prevent transmission of the disease (*3*). In addition, simple chemistry and resulting low manufacturing costs are important requirements to ensure access to new medicines for patients in low income countries where malaria is endemic.

Pantothenate (pantothenic acid, vitamin B5, see Figure 1A) is the only water-soluble vitamin that is essential for malaria parasite viability (*4*). A variety of pantothenate analogues, including pantoyltaurine, substituted pantoyltaurylamides, sulphonamides, and pantothenones, have shown antimalarial activity in avian malaria models (*5*). Pantetheine analogues based on N1-(substituted) pantothenamides were also shown to exert antimicrobial activity (*6*, *7*). In *E. coli* and *S. aureus*, pantothenamides were reported to enter the coenzyme A (CoA) biosynthesis pathway following phosphorylation by pantothenate kinase (PANK) and form antimetabolites that block CoA and acyl-carrier dependent processes (*7*, *8*).

**Figure 1.**
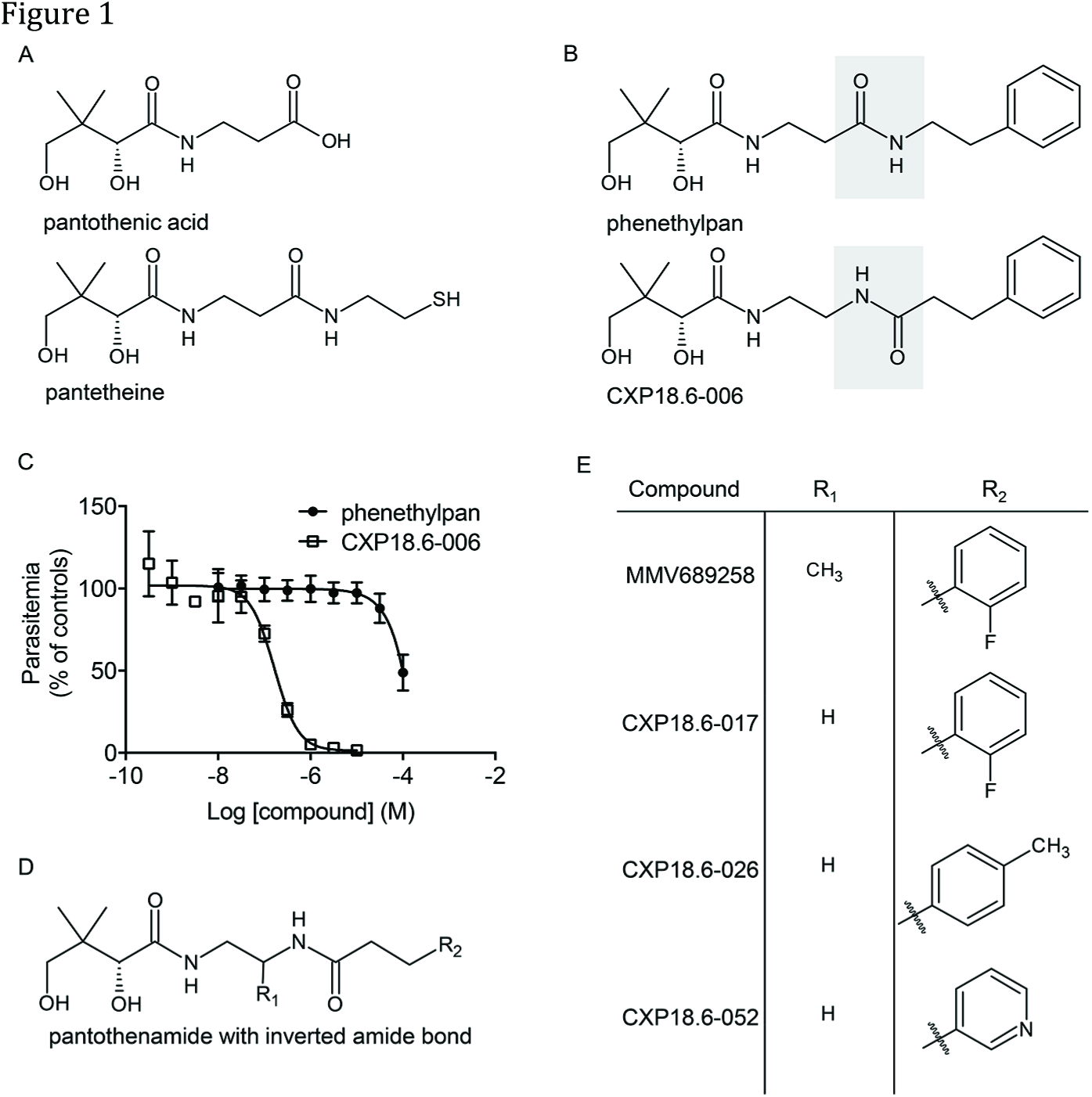
Generation of stable pantothenamides. A) Structures of pantothenic acid, the starting point of CoA biosynthesis, and panthetheine, a natural pantothenamide that is the substrate of vanins. B) Structure of phenethylpan, starting point for chemical optimization and its stabilized analog CXP18.6-006. C) Antimalarial activities of phenethylpan and CXP18.6-006 when tested against *P. falciparum* NF54 asexual blood-stage parasites. The figure shows mean parasite density relative to control. Error bars indicate standard deviations determined from three (phenethylpan) or two (CXP18.6-017) independent replicates. D) Core structure of pantothenamides described in this paper. E) Structures of lead pantothenamide compounds.

In spite of their potent antimicrobial action *in vitro*, pantothenate and pantetheine derivatives (Figure 1A) have never been developed into clinically effective drugs. This is due to their intrinsic instability in body fluids as they are hydrolyzed by the ubiquitous pantetheinase enzymes of the vanin family (*9*). Previous work has shown that classical pantothenamides are active against *P. falciparum in vitro*, provided that plasma pantetheinase activity is inhibited by heat inactivation or addition of a small molecule vanin inhibitor (*9*, *10*). Recent reports have shown that structural modifications of pantothenamides will increase their stability in serum, *in vitro* (*11*, *12*). Studies on the mode of action of antimalarial pantothenamides are conflicting as both formation of antimetabolites and inhibition of PANK have been reported (*13*-*15*). In rodent malaria parasites, the pantothenate transporter and the downstream enzymes phosphopantothenylcysteine synthase and phosphopantothenylcysteine decarboxylase are essential for development of the parasite in the mosquito midgut, suggesting that a drug interfering with this pathway may prevent disease transmission (*16*-*18*). These findings have motivated our search for a new class of pantetheinase-resistant pantothenamide antimalarials. We present a new class of pantothenamide analogues that are both stable, with the labile amide bond replaced by a pantetheinase-resistant bioisostere, and potent, with low nanomolar activity towards asexual and sexual blood stages of *P. falciparum*. Targeted metabolomics revealed that pantothenamides are metabolized by CoA biosynthetic enzymes, leading to antimetabolites that suppress acetyl-CoA synthesis. We further demonstrate that these compounds are active *in vivo*, using a humanized rodent infection model, and demonstrate that acetyl-CoA synthetase and acyl-CoA synthetase 11 play crucial roles in the sensitivity of parasites to pantothenamide compounds.

## Results

### Development of stable pantothenamide bioisosteres

The starting point for chemical optimization of a new class of pantothenamides was phenethylpan (Figure 1B), a compound with potent antimalarial activity when protected from degradation by serum vanins through heat-inactivation (*10*) or addition of a vanin inhibitor (*19*). Inversion of the amide yielded a pantothenamide bioisostere CXP18.6-006 (Figure 1B) that was completely resistant to hydrolysis by serum-derived vanins as determined *in vitro* and *in vivo* (Figure S1). As a result, the compound was found to be several orders of magnitude more potent than the parent compound phenethylpan when tested for its ability to block replication of asexual blood stages of *P. falciparum* NF54 parasites in serum-containing medium (Figure 1C). The structure-activity relationship in the pantothenamide series was further explored by a number of modifications at various positions of the molecule (Figure 1D). Key findings were that 1) the two hydroxyls and the two methyl groups are essential for activity, 2) an ethyl provides the optimal chain length for both carbon linkers, 3) alkyl substitution with (S)-methyl at R1 yielded a 10-fold increase in potency, 4) the *R* configuration of the secondary alcohol and the *S* configuration of the methyl at R1 are preferred and 5) substituted aryl, alkyl and hetero-aromatic side-chains are tolerated at position R2. From this series of substitutions, four primary compounds (Figure 1E) emerged that provided potent (nM) antimalarial activity (Table S1).

### Pantothenamides block asexual blood stage replication and parasite transmission

We investigated the effects of our four primary compounds at various stages of the parasite life cycle (Figure 2A). The compounds inhibited replication of NF54 asexual blood-stage parasites with IC_50_ values < 110 nM, with MMV689258 being the most potent compound at an IC_50_ of 5 nM (Figure 2B and Table S1), while its *SR* enantiomer MMV884968 was inactive (Table S1). MMV689258 was active against a diverse panel of *P. falciparum* strains with resistances against a wide range of marketed antimalarial drugs, suggesting a novel mechanism of action (Figure 2C and Table S2). Despite highly potent activity against blood-stage parasites, only weak to moderate activity was seen against developing intra-hepatocytic schizonts (Figure 2D and Table S1). Based on the observation that a number of enzymes in the pantothenate-CoA biosynthesis route are essential for parasite development in the mosquito midgut (*16-18*), we also investigated pantothenamide effects against the transmission stages of the parasite. To this end, mature gametocytes were incubated with compound for 24 hours and fed to *Anopheles stephensi* mosquitoes in a standard membrane feeding assay (SMFA) (*20*). Eight days after feeding the infection status of the mosquitoes was analyzed. The results show that both CXP18.6-017 and MMV689258 prevent mosquito infection, with IC_50_ values that are comparable to their activity against asexual blood stages (Figure 2E and Table S1). This transmission-blocking activity relies on a gametocytocidal mode of action, as indicated by gametocyte viability assays that revealed IC_50_ values in line with the transmission blocking activity (Figure 2F and Table S1).

**Figure 2.**
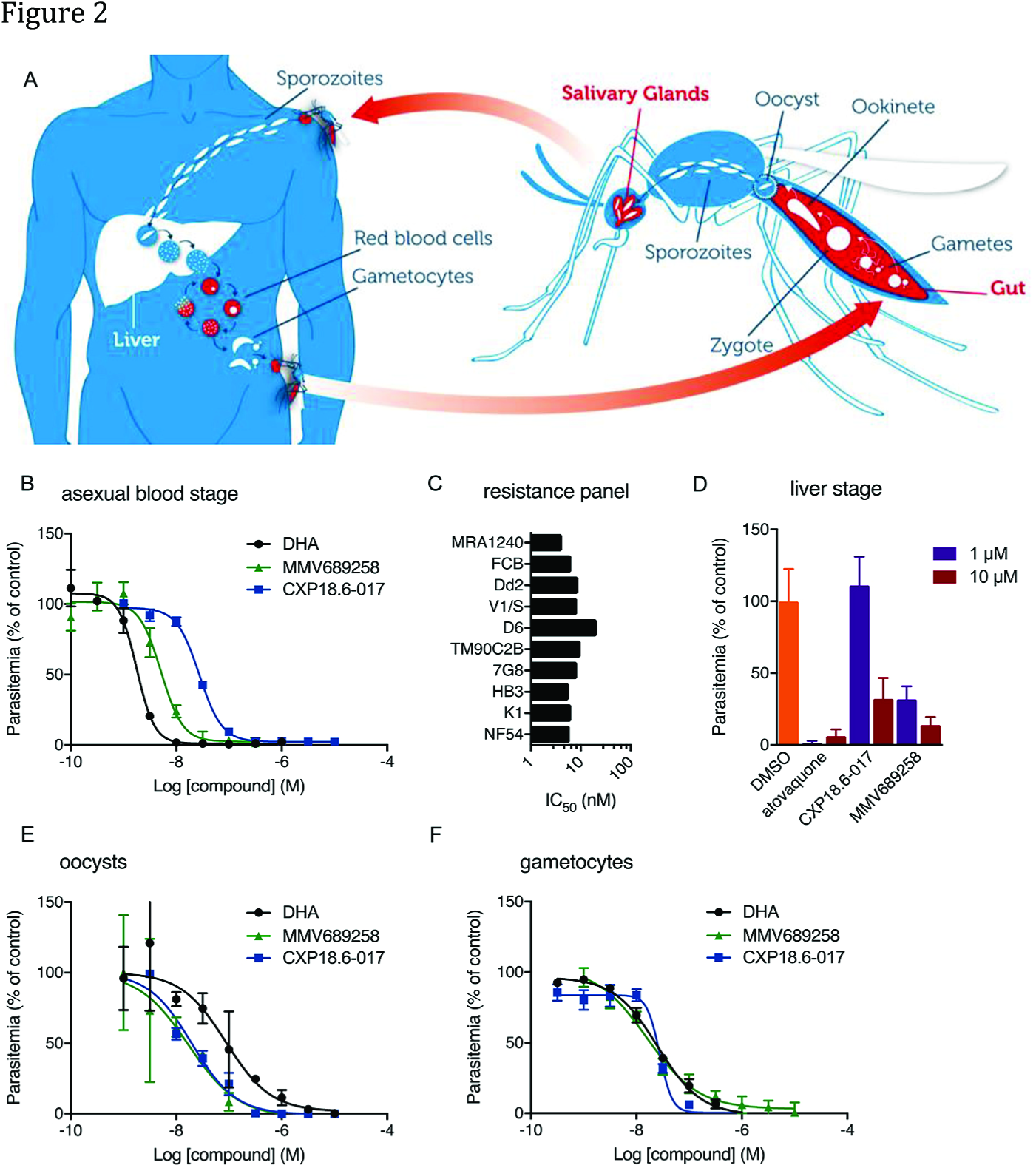
*In vitro* antimalarial activities of pantothenamides. A) Life cycle of the human malaria parasite indicating the key parasite stages in the human and mosquito host. B) Activity of pantothenamides against *P. falciparum* NF54 asexual blood-stage parasites. C) Activity of MMV689258 against a panel of drug resistant *P. falciparum* strains and, for comparison, drug sensitive NF54 parasites. D) Analysis of compound effects on the development of *P. falciparum* NF54 in human primary liver cells. All panels show mean values and standard deviations for 2-3 independent experiments. E) Transmission blocking activity. Mature *P. falciparum* stage V gametocytes from luminescent reporter strain NF54-HGL were exposed to compound for 24 hours prior to feeding to *A. stephensi* mosquitoes. Eight days post feeding, the infection status of the mosquitoes was assessed by luminescence analyses. F) Gametocytocidal activity against *P. falciparum* NF54 gametocytes harvested at day 11 post induction of gametocytogenesis.

### Pantothenamides are converted to Co-A conjugates that block acetyl-CoA synthesis

For all pantothenamides tested, the activity against asexual blood-stage parasites could be outcompeted with an excess of pantothenate (Figure S2). Given this competitive mode of action, we addressed whether the compounds interfere with PANK, which has been shown to be the rate-limiting step in the pantothenate to CoA conversion in other organisms (*21*). Phosphorylation of pantothenate was detected both in lysates of infected red blood cells and in uninfected red blood cells (Figure S3A). In order to investigate the presence of a genuine PANK enzyme in parasites we produced a baculovirus-expressed recombinant form of the predicted *P. falciparum* pantothenate kinase 1 (PfPANK1). This protein was inactive *in vitro* (data not shown). However, an antiserum generated against this protein immunoprecipitated PANK activity from *P. falciparum* lysates (Figure S3B and S3C). Together with recently published genetic data (*15*), this suggests that PfPANK1 represents a genuine PANK enzyme. Subsequent *in vitro* assays revealed that the pantothenamides are relatively poor inhibitors of human and parasite PANK activity, and the resulting IC_50_ values do not align with the *in vitro* antimalarial activities (Figure S3D and Table S1). This suggests that parasite PANK activity is not the primary target for these compounds.

Using an established LC-MS-based metabolomics methodology for predicting the mode of action of antimalarial compounds (*22*), we tested the parasite response to pantothenamides using purified trophozoite-stage parasites. *In vivo* CoA modifications of pantothenamides have previously been reported in parasites and bacteria (*23*). Therefore, we predicted possible chemical derivatives based on the structural and chemical features of the pantothenamides (Table S3) as well as the endogenous metabolites involved in cellular CoA production (Figure 3A). HPLC-based metabolomics confirmed that all of the tested antimalarial pantothenamides were ultimately converted to pantothenamide-CoA-conjugates in red blood cells containing trophozoite stage parasites (Figure 3B and Figure S4A). Similarly, as expected due to the gametocytocidal activity of the compounds, the CoA-conjugated form was also detected in the gametocyte stage of the parasite (Figure S4B). In comparison with CoA, that has a terminal thiol, the aryl group (R_2_, Figure 1C) of the pantothenamides blocks acetylation by acetyl-CoA synthetase. Indeed, we did not find pantothenamide metabolites downstream of acetyl-CoA synthetase, confirming that the conversion of pantothenamides terminates at this enzyme.

**Figure 3.**
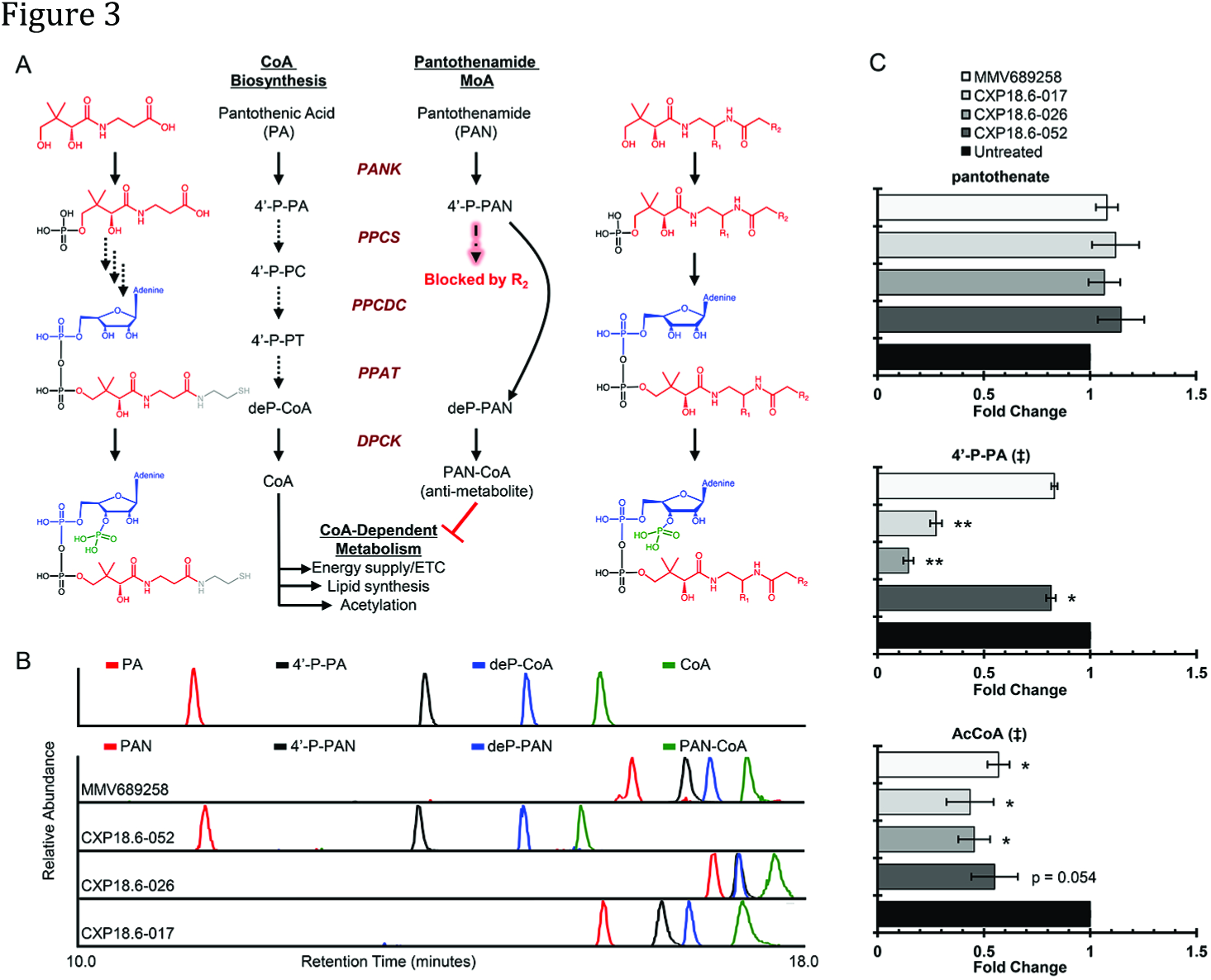
CoA biosynthesis and pantothenamide metabolism in MACS-purified trophozoites. A) Schematic for the endogenous pantothenate/CoA biosynthesis pathway (left) or pantothenamide metabolism (right). Structures are displayed for the metabolites of interest and colored to denote key functional residues. Enzyme abbreviations are displayed in red between the metabolic routes to denote the utilization of these enzymes in both pathways. Enzymatic blockages in the pantothenamide route are denoted within in the pathway in red. Abbreviations are: PA = pantothenic acid, 4′-P-PA = 4′-phosphopantothenate, 4′-P-PC = 4′-phosphopantothenoyl-L-cysteine, 4′-P-PT = pantetheine-4′-phosphate, deP-CoA = dephosphoCoA, CoA = Coenzyme-A, PAN = pantothenamide, 4′-P-PAN = 4′-phosphopantothenamide conjugate, deP-PAN = dephosphoCoA pantothenamide conjugate, PAN-CoA = pantothenamide CoA conjugate, PANK = pantothenate kinase, PPCS = phosphopantothenate-cysteine ligase, PPCDC = phosphopantothenoylcysteine decarboxylase, PPAT = pantetheine-phosphate adenyltransferase, DPCK = dephospho-CoA kinase. B) Representative extracted ion chromatograms (EICs) for pantothenate and panothenamide metabolism. The upper panel displays peaks for endogenous compounds run as pure standards (1μM), while antimetabolite peaks are from cellular extracts. The x-axis denotes the retention time of the detected metabolite (colors match schematic in panel A) and the y-axis denotes the relative abundance (100% per respective EIC). All peaks identified were unique to the drug treatment (untreated EICs not shown for clarity). C) Endogenous metabolic alterations in trophozoite stage parasites treated with 10XIC_50_ compound for 2.5 hours. Y-axis denotes the pantothenamide tested and the x-axis is the average fold change (±SE) relative to a paired untreated control. Each sample was collected in technical triplicate for n = 3 biological replicates. ‡ in the title denotes statistical significance at p < 0.05 by one-way ANOVA between all groups. * p < 0.05, ** p < 0.01, paired t-tests versus untreated.

Targeted analyses of endogenous metabolites in the CoA pathway indicated that acetyl-CoA levels in asexual blood-stage infected erythrocytes were significantly lower in the presence of the pantothenamides (Figure 3C, lower graph). Two of the compounds, CXP18.6-017 and CXP18.6-26, also resulted in large decreases of 4′-phosphopantothenate (Figure 3C, middle graph), a depletion that most likely results from competition with endogeneous pantothenate. Indeed, the detection of the phosphorylated form of the pantothenamides confirms that they are a substrate for PANK and may compete with pantothenate when binding to the catalytic site. In gametocytes, incubation with 1 μM MMV689258 for 2.5 hours resulted in similar changes in levels of endogenous metabolites in the CoA biosynthesis pathway (Figure S4C). No additional significant changes were identified in asexual parasite metabolism (Figure 3C, upper graph and Figure S4D), further demonstrating CoA pathway specificity.

In order to delineate red blood cell and parasite-mediated metabolism, saponin lysis was used to isolate trophozoite-stage parasites from infected human RBCs. Incubation of free parasites with 1 μM MMV689258 for three hours in pantothenate-free media, led to accumulation of high levels of the phospho-and dephospho-CoA forms of the compound and lower levels of the parent compound and final CoA-conjugated form (Figure S5A). In control experiments with uninfected red blood cells we also observed the generation of MMV689258-antimetabolites, in particular the levels of MMV689258-CoA were comparable to those observed in parasites (Figure S5A). This was quite unexpected since a parallel experiment performed with isotopically labeled pantothenate demonstrated that red blood cells do not readily metabolize pantothenate. Compared to levels observed in free parasites, *de novo* generated labeled acetyl-CoA in red blood cells was negligible, as values were at or below background noise (Figure S5B). Although human red blood cells lack de novo CoA synthesis (*24*), they do contain hPANK2 and Coenzyme-A synthase (COASY) which explains the generation of the MMV689258-CoA conjugate in red blood cells (Figure 3A). In agreement with the lack of *de novo* synthesis of CoA in red blood cells, the pantothenamide did not affect acetyl-CoA levels in these cells (Figure S5C). In contrast to red blood cells, parasite acetyl-CoA levels dropped fifteenfold following treatment with MMV689258 (Figure S5C), indicating that the reduction in acetyl-CoA levels observed in infected erythrocytes is due to a reduction of parasitic rather than red blood cell acetyl-CoA.

The observation that uninfected red blood cells convert the pantothenamide bioisosteres to the CoA-conjugated form, prompted us to investigate the effect of pre-existing drug-conjugates on parasite infection. Red blood cells were pulsed with compound for 3 hours, followed by thorough washing and a 24-hour chase to allow conversion of the parent compound to the drug-conjugates and presumed antimetabolite. Subsequently, the cells were infected with synchronized parasites that were allowed to proliferate for 72 hours. In this protocol, washout of the reference compound dihydroartemisinin did not inhibit parasite growth in drug-exposed cells (Figure S6A). In contrast, a 3 hour exposure of red blood cells to chloroquine, which is known to accumulate in red blood cells (*25*), was sufficient to exert a reduction in parasite replication. Likewise, parasites replicated poorly in red blood cells exposed to a pulse of MMV689258 prior to infection, suggesting that red blood cell derived antimetabolites contribute to the pantothenamide mode of action. Parallel metabolomics experiments indicated that following an exposure of red blood cells to MMV689258 for 3 hours, the upstream products rapidly diminish and the resulting pantothenamide-CoA conjugate (PAN-CoA) persists for at least 72 hours (Figure S6B).

### Mutations in acetyl-CoA synthetase and acyl-CoA synthetase 11 confer resistance to pantothenamides

To further elucidate the molecular mechanism of action of the pantothenamides, resistant parasites were selected by increased exposure to CXP18.6-052. Selected clones showed IC_50_ values that were 50 to 70 fold higher compared to the activity against wild-type parasites and were cross-resistant to other pantothenamides tested, suggesting a shared mechanism (Figure 4A). Whole genome sequencing of five clones from two independent flasks revealed that all clones had mutations in the fatty acid anabolism protein acyl-CoA synthetase 11 (ACS11) (PF3D7_1238800). For two clones, the mutation results in an E660K amino acid change while three clones contained a K462N substitution. In *Plasmodium*, the ACS family of enzymes is an expanded 13-member gene family (*26*) and sequence alignments clearly define highly conserved amino acids in the active site of the enzyme, specifically residues that bind CoA. A common mutation was found in acetyl-CoA synthetase (AcCS) (PF3D7_0627800) resulting in a T627A amino acid change (Figure 4B). While this residue is not conserved across species, it does fall within the CoA binding pocket (Figure 4C), suggesting it leads to activity or specificity changes in the enzyme. The inhibition of AcCS logically explains the observed decrease in cellular acetyl-CoA levels following pantothenamide treatment, particularly because acetyl-CoA synthetase has been shown to be a major contributor to cellular acetyl-CoA levels in *P. falciparum* (*27*). All clones also showed common mutations in a single *var* gene (PF3D7_0900100), which encodes a highly polymorphic erythrocyte adhesion protein and is unlikely to contribute to the drug-resistant phenotype we observed.

**Figure 4.**
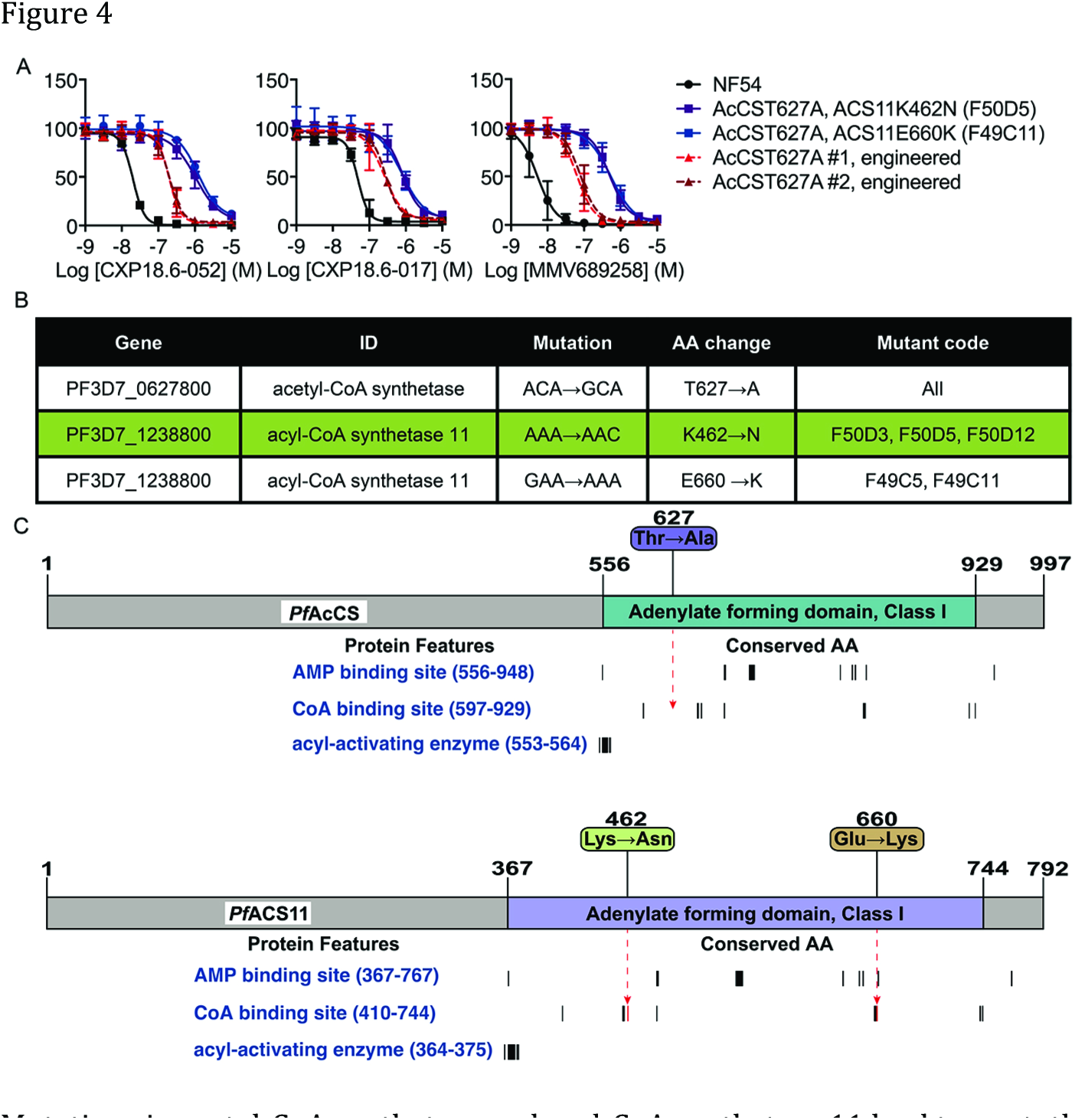
Mutations in acetyl-CoA synthetase and acyl-CoA synthetase 11 lead to pantothenamide resistant parasites. A) Drug sensitivity assays with *P. falciparum* NF54 parasites selected for resistance against CXP18.6-052. The figure shows average values relative to control from asexual blood stage replication assays for two independently selected lines and, for comparison, NF54 wildtype parasites. Error bars indicate standard deviations determined from two independent replicates. B) Table of mutations found in resistant parasite lines that result in protein coding amino acid changes. These mutations were identified from whole genome sequencing, by comparison to the 3D7 reference genome and wild-type NF54 parental line. Multiple clones were sequenced for confirmation from each independently derived line. C) Schematic of protein domains and mutations. NCBI protein sequences were used to map protein domains annotated by the Conserved Domain Database. Black vertical bars denote conserved amino acids within a protein feature. Arrows demonstrate the mutation position within the protein features and mutations that occur in conserved amino acids are denoted as red.

To investigate the causal role of the AcCS T627A mutation in the drug resistance phenotype, this mutation was engineered in the *P. falciparum* NF54 background using the CRISPR/Cas9 system. In comparison with parasites that contained both an AcCS and ACS11 mutation, the AcCS T627A parasites showed an intermediate resistance phenotype (Figure 4A). Although resistant parasites were able to produce gametocytes that infected *Anopheles stephensi* mosquitoes (Figure S7), asexual competition assays demonstrated that blood stages have a fitness cost and a dramatically reduced ability of the mutant alleles to spread in the parasite population (Figure S8)

### MMV689258 suppresses parasitemia in a model for human malaria infection

Pharmacokinetic experiments with MMV689258 revealed moderate clearance of 50 ml/min/kg in mice and 23ml/min/kg in rats with elimination half-lives of 1.8 and 3.6 hours, respectively (Figure S9 and Table S4). The volume of distribution was 2.52 l/kg in mice and 1.71l/kg in rats and the compound was well absorbed upon oral dosing (bioavailability of 33.6% in mice and 63.4% in rats). Assessment of compound levels in urine and bile collected from rats showed that renal clearance contributed 26% to total clearance, whereas biliary clearance was negligible (Tables S4 and S5). Based on observed intrinsic clearance in rat hepatocytes of 6.6 μl/min/k (Table S6), predicted *in vivo* hepatic clearance would amount to 14 ml/min/kg, which is well line with the non-renal portion (17 ml/min/kg) of the total observed *in vivo* clearance suggesting that the main route of elimination is through the liver.

To study the relationship between blood levels and parasite clearance, MMV689258 was tested in a humanized mouse model for *P. falciparum* infection. Based on the relatively short half-life in mice, the compound was initially dosed once daily for four days. At day 7 after infection, parasitemia was fully cleared (>99.9% activity compared to untreated control mice) by four daily doses of 50 mg/kg and higher, whereas the daily 25 mg/kg dose reduced parasitemia by 99.8% (Figure 5A). At the higher dosages, the rate of parasite clearance was similar to that observed for chloroquine. Blood concentrations of MMV689258 in humanized mice showed a similar profile as observed in initial mouse pharmacokinetic experiments (Figure 5C). Analyses of dose-normalized data indicated that C_max_ and AUC_0-24_ values were super-proportional with dose, suggesting possible saturation of clearance (Figure S10). With measured fractional mouse plasma protein binding of 32.8% (Table S6), total blood levels of 48.6 ng/ml are required to achieve a free fraction above the *in vitro* determined IC_99_ (45 nM) of MMV689258. For all dosages, blood levels dropped below this level within 24 hours. To further investigate the exposure-efficacy relationship, MMV689258 was administered as a single dose in the humanized mouse model. At doses from 25 to 200 mg/kg at day 3 after infection, parasitemia was reduced by 77-99.9% at day 7 compared to untreated control mice (Figure 5B). The combined data indicate that a relatively short exposure above the IC99 level is sufficient to achieve efficacious clearance of asexual blood-stage parasites. The accumulation of the MMV689258-CoA-conjugate as observed in red blood cells (Figures S5A and S6B) may contribute to this seemingly prolonged duration of compound action.

**Figure 5.**
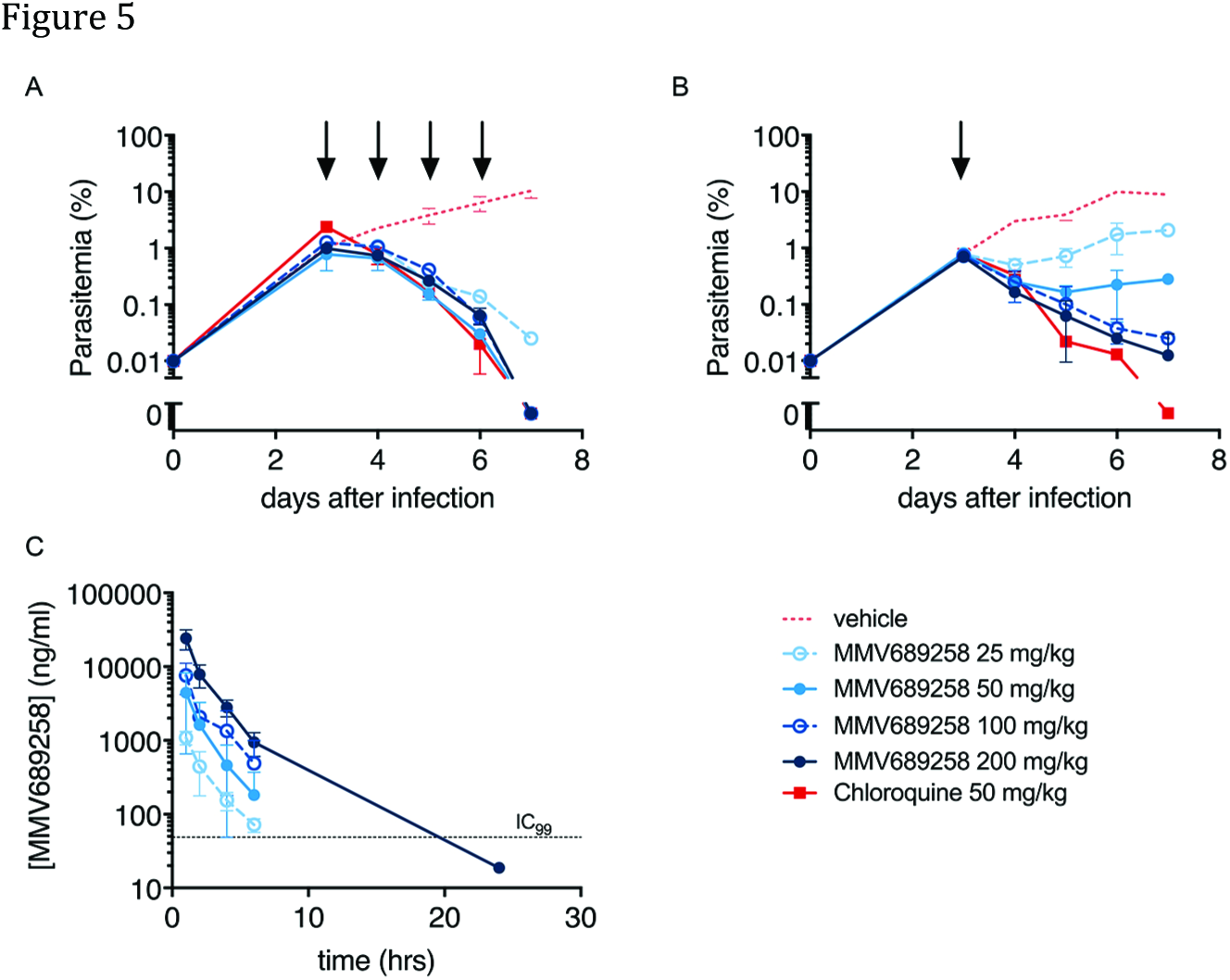
*In vivo* efficacy and pharmacokinetics of MMV689258. Female NODscidIL2Rγ^null^ mice were engrafted with human red blood cells and infected with *P. falciparum* at day 0. At day 3 post infection, mice were dosed with the compounds at doses indicated in the figure legend by oral gavage. The figures show average values and standard deviations from two mice per drug/dose combination. A) Parasitemia in time following daily doses at four consecutive days (black arrows). B) Parasitemia in time following a single dose of compound (black arrow). C) Plasma levels of MMV689258 as a function of time following the first administration of compound at the dose indicated in the figure legend.

In all animal experiments, drug administration was well tolerated and no obvious acute adverse effects were noted. In keeping with this apparent lack of toxicity, none of the pantothenamides tested showed cytototoxic activity against HepG2, HC04 cell lines or human primary hepatocytes (Table S1). Metabolic profiling in primary human hepatocytes lines showed that only a small fraction of the total pool of compound was converted to phospho-and dephospho-CoA conjugates, whereas the CoA conjugate was not detected (Figure S11A). Levels of endogenous metabolites in the pantothenate-CoA pathway were unaltered in pantothenamide-exposed primary hepatocytes (Figure S11B). Additionally, MMV689258 and CXP18.6-017 did not inhibit hepatic CYP450 enzymes at concentrations up to 20 μM, indicating a low risk for drug-drug interaction (Table S1). Selection of a pantothenamide clinical candidate will require further analyses of safety in *in vitro* and *in vivo* models.

## Discussion

Here we describe a new, stable class of pantothenamide bioisosteres that display potent activity against asexual and sexual blood stages of *P. falciparum* and exert antimalarial activity in a humanized mouse model for *P. falciparum* infection. Our findings strongly support the advancement of stable pantothenamide bioisosteres for further evaluation of efficacy and safety in preclinical models and humans. These compounds constitute an important addition to the antimalarial portfolio, as they 1) are straightforward to synthesize and are expected to have low costs of good, 2) are transmission-blocking, and 3) target a biochemical pathway that is not covered by currently marketed drugs.

Previous studies in bacteria have demonstrated that antibiotics based on the pantothenate/pantetheine scaffold are labile in biological fluids due to hydrolysis of the amide bond by serum vanins (*9*). Our data reveals that inversion of the amide bond afforded a considerable increase of stability whilst preserving the biological activity. Antimalarial activity was further improved by modification of the carbon linker and the aromatic ring of the side chain. Metabolomic analyses indicated that the pantothenamides utilize the CoA biosynthetic enzymes of both parasites and red blood cells and are ultimately converted into CoA-drug conjugates. As the pantothenamides lack the pantothenate carboxyl group they bypass PPCS and PPCDC and are converted to the CoA-conjugate by PPAT and DPCK upon phosphorylation by PANK. Consistent with this mechanism, our medicinal chemistry efforts revealed a narrow pharmacophore. All substitutions of functional groups that are essential for processing by PANK, PPAT and DPCK, including the two alcohols and the dimethyl motif, led to a loss of antimalarial activity. The mechanism described here is consistent with previous observations of CoA conjugation of pantothenate analogues in infected red blood cells (*15*).

Interestingly, uninfected red blood cells pre-treated with a pantothenamide demonstrated a prophylactic capacity by significantly reducing subsequent proliferation of a naïve parasite population. Metabolomics revealed that the resulting CoA-conjugate is formed rapidly and is stably retained within the uninfected cells after removal of the parent compound. This unprecedented host cell activity could also explain the apparent disconnect between the observed plasma levels versus *in vivo* efficacy in the mouse model; an idea backed by the rapid clearance of parasites after a single treatment. Further development of a pantothenamide clinical candidate and prediction of the efficacious human dose would benefit from a better understanding of the exposure-effect relationship, which is the subject of our current studies.

We propose that conjugation of the pantothenamides by CoA biosynthetic enzymes produces antimetabolite(s) that interfere with downstream CoA-dependent processes. The observed decrease of acetyl-CoA levels in drug exposed parasites is consistent with such a mechanism. Furthermore, selection and sequencing of drug resistant parasite lines revealed mutations in the CoA-binding enzymes AcCS and ACS11. Mutations in these active site residues may lead to altered enzyme specificity, thereby excluding the antimetabolite from altering its activity, albeit this comes at an observed parasite fitness cost. We are aware that the generation of antimetabolites and interference with CoA-dependent processes in host cells is clearly a potential source of unwanted side effects. However, we did not observe a significant level of antimetabolite formation in primary hepatocytes, nor did we observe acute toxic effects or adverse events in rats or mice upon administration of oral doses up to 200 mg/kg, suggesting a specificity for the parasite enzymes over the host.

In conclusion, our study identifies stable pantothenamide bioisosteres as a new class of antimalarial compounds with a unique mode of action. These compounds are equally active against the parasite stages that cause clinical disease and those that drive onward transmission by the mosquito, and provide important starting points for the development of next-generation malaria-eliminating drugs.

## Methods

### Chemistry

Unless noted otherwise, materials were purchased from commercial suppliers and used as received. All air and moisture sensitive reactions were carried out under an inert atmosphere of dry nitrogen. DCM was dried over Na2SO_4_ prior to use. Column chromatography was performed using Acros silica gel (0.035-0.070 mm, 6 nm). NMR spectra were recorded at 298 K on a Varian 400 (400 MHz) spectrometer in the solvent indicated. Chemical shifts are given in parts per million (ppm) with respect to tetramethylsilane (0.00 ppm), or CHD_2_OD (3.31 ppm) as internal standard for 1H NMR. Coupling constants are reported as/ values in hertz (Hz).

### Synthesis ofCXP18.6-006, CXP18.6-017, CXP18.6-026

General procedure A: to a solution of carboxylic acid **B** (0.5 mmol) in MeCN/H_2_O (30:1, 4.3 mL) were added HOBt (0.6 mmol), NaHCO3 (0.6 mmol), EDCI (0.6 mmol) and a solution of amine **A** (for the synthesis see ref 2, 0.6 mmol) in MeCN/H2O (0.7 mL). The progress of the reaction was monitored using LC-MS and upon completion, the reaction was quenched by the addition of saturated aqueous NH_4_Cl solution (15 mL) and the mixture was extracted twice using EtOAc (15 mL). The combined organic layers were dried over Na2SO_4_ and filtered before concentration under reduced pressure. The residue was purified by flash column chromtography (DCM/MeOH = 98:2 → 80:20) to afford the product.

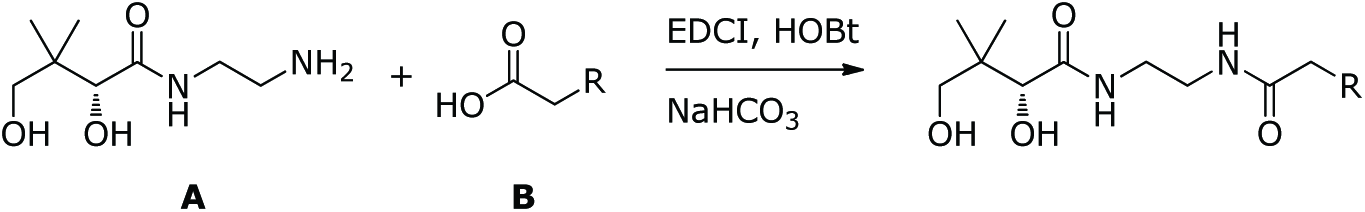

#### CXP18.6-006

According to general procedure A. Yield: 51%, pale yellow oil. 1H NMR (400 MHz, CD_3_OD): *δ* 7.30-7.15 (m, 5H), 3.88 (s, 1H), 3.46 (d, *J* = 11.0 Hz, 1H), 3.36 (d, *J* = 11.0 Hz, 1H), 3.31-3.21 (m, 4H), 2.93-2.87 (m, 2H), 2.50-2.44 (m, 2H), 0.92 (s, 3H), 0.92 (s, 3H).

#### CXP18.6-017

According to general procedure A. Yield: 35%, colorless oil. 1H NMR (400 MHz, CD3OD): *δ* 7.28-7.17 (m, 2H), 7.11-6.98 (m, 2H), 3.88 (s, 1H), 3.46 (d, *J* = 11.0 Hz, 1H), 3.39 (d, *J* = 11.0 Hz, 1H), 3.33-3.22 (m, 4H), 2.98-2.92 (m, 2H), 2.51-2.54 (m, 2H), 0.92 (s, 3H), 0.92 (s, 3H).

#### CXP18.6-026

According to general procedure A. Yield: 56%, colorless oil. 1H NMR (400 MHz, CD_3_OD): *δ* 7.08 (app s, 4H), 3.88 (s, 1H), 3.46 (d, *J* = 11.0 Hz, 1H), 3.39 (d, *J* = 11.0 Hz, 1H), 3.33-3.21 (m, 4H), 2.88-2.82 (m, 2H), 2.47-2.40 (m, 2H), 2.28 (s, 3H), 0.92 (s, 3H), 0.92 (s, 3H).

### Synthesis ofCXP18.6-052

General procedure B: to a suspension of carboxylic acid B (0.72 mmol) in MeCN/H_2_O (19:1, 4 mL) was added EDCI (0.79 mmol). After stirring for 10 min an almost clear solution was obtained. Amine A (for the synthesis see ref 2, 0.79 mmol) was added, followed by DIPEA (0.79 mmol). The progress of the reaction was monitored using LC-MS and upon completion, silica gel was added. The mixture was then concentrated under reduced pressure and purified by flash column chromatography (MeCN/MeOH = 4:1 → 2:1) to afford the product.

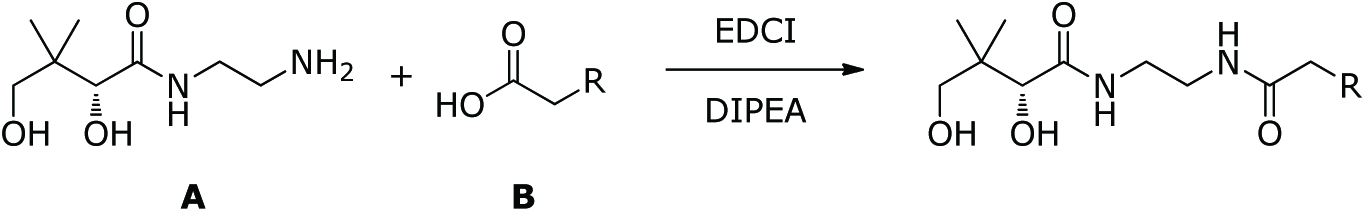

#### CXP18.6-052

According to general procedure B. Yield: 30%, colorless oil. ^1^H NMR (400 MHz, CD3OD): *δ* 8.42 (d, *J* = 1.8 Hz, 1H), 8.37 (dd, *J* = 4.9,1.6 Hz, 1H), 7.72 (ddd, *J* = 7.9, 2.2,1.7 Hz, 1H), 7.37 (ddd, *J* = 7.8, 4.9, 0.8 Hz, 1H), 3.88 (s, 1H), 3.46 (d, *J* = 11.0 Hz, 1H), 3.39 (d, *J* = 11.0 Hz, 1H), 3.30-3.20 (m, 4H), 2.96 (t, *J* = 7.5 Hz, 2H), 2.51 (t, *J* = 7.5 Hz, 2H), 0.92 (s, 3H), 0.91 (s, 3H).

### Synthesis of MMV689258

To a stirred solution of ((*S*)-2-Hydroxy-1-methyl-ethyl)-carbamic acid tert-butyl ester **1** (2.5 g, 14.26 mmol) in dry THF (28 ml) were added phthalimide **2** (2.3g, 15.69 mmol) and PPh3 (4.11g, 15.69 mmol). DEAD (2.73 g, 15.69 mmol) was then added dropwise to the stirred solution at room temperature and maintained for 16h. The reaction mixture was then concentrated under reduced pressure and the residue was purified by column chromatography (70:30 −50:50 hexanes-EtOAc) to afford compound **3** (4 g, 92%) as white solid.

To a stirred solution of compound **3** (2.0 g, 6.57 mmol) in dioxane (20 ml) was added 4M dioxane-HCl (20 ml) at 0°C. It was stirred at RT for 6h. It was concentrated under reduced pressure to afford compound **4** (1.5 g, 95%) as white solid.

To a stirred solution of compound **5** (180 mg, 1.07 mmol) in THF (10 ml) were added Et3N (0.747 ml, 5.36 mmol), HATU (611 mg, 1.61 mmol) and compound **4** (515 mg, 2.14 mmol). It was stirred at RT for 16h. TLC (30% EtOAc-Hexane) showed completion of the reaction. It was diluted with water (20 ml) and extracted with EtOAc (50 ml), washed with brine (10 ml), dried over Na2SO4 and concentrated under reduced pressure. It was purified by combiflash (30% EtOAc-Hexane) to afford compound **6** (330 mg, 87%) as off white solid.

To a stirred solution of compound **6** (330 mg, 0.93 mmol) in EtOH (20 ml) was added hydrazine hydrate (745 mg, 14.90 mmol). It was heated at 50°C for 2h. It was cooled to RT, filtered and concentrated under reduced pressure. The resulting residue was suspended in Et2O (20 ml) and filtered, washing thoroughly with Et2O (20 ml). The combined filtrates were concentrated under reduced pressure to afford compound **7** (150 mg, 72 %) as gum.

To a stirred solution of compound **8** (75 mg, 0.32 mmol) in THF (20 ml) were added Et3N (0.221ml, 1.59 mmol), HATU (181.25 mg, 0.48 mmol) and compound **7** (142.37 mg, 0.64 mmol). It was stirred at room temp for 16h. It was diluted with water (20 ml) and extracted with EtOAc (50 ml), washed with saturated NaHCO3 (20 ml), brine (10 ml), dried over Na2SO_4_ and concentrated under reduced pressure. It was purified by column chromatography (3% MeOH-DCM) to afford compound **9** (110 mg, 78%) as gum.

Compound **9** (90 mg, 0.203 mmol) was taken in MeOH (10 ml) and was degassed with argon for 10 minutes. Pd/C (50 mg, 10% moist) was added and the mixture was subjected to hydrogenation in a Parr vessel at 50psi for 16h. It was filtered through celite, concentrated under reduced pressure and purified by preparative TLC (4% MeOH-DCM) to afford MMV689258 (40 mg, 55 %) as colourless gum. 1H NMR (400 MHz, DMSO-*d*_6_) **8** 0.79 (ds, 6 H), 0.95 (d, 3H), 2.32 (t, 2H), 2.81 (t, 2H), 2.99-3.02 (m, 1H), 3.12-3.18 (m, 2H), 3.27-3.32 (m, 1H), 3.72 (d, 1H), 3.83-3.86 (m, 1H), 4.46 (t, 1H, −OH), 5.38 (d, 1H, −OH), 7.08-7.14 (m, 2H), 7.21-7.28 (m, 2H), 7.68-7.72 (m, 2H).

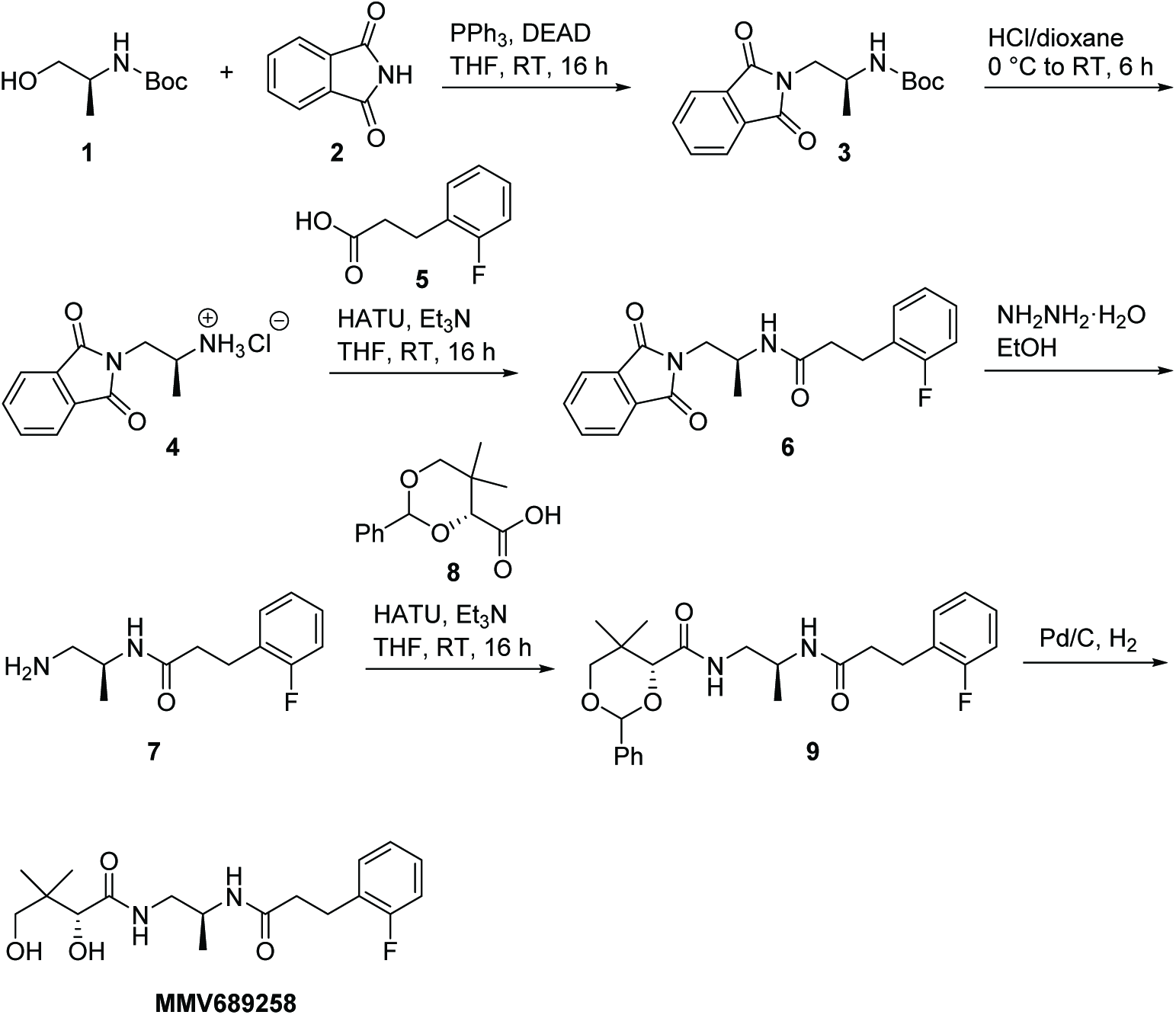

### *In vitro* parasitology

*Plasmodium falciparum* strain NF54 was cultured in RPMI1640 medium supplemented with 25 mM NaHCO3,10% human type A serum and 5% (v/v) human type 0 red blood cells (Sanquin, the Netherlands) in a semi-automated culture system (*28*). Replication assays for asexual blood-stage parasites were performed by diluting parasites in RPMI1640 medium with 10% human serum to a parasitemia of 0.83% and a hematocrit of 3%. 30 μl of diluted parasites were combined with 30 μl of compound serially diluted in DMSO and RPMI1640 medium to reach a final DMSO concentration of 0.1% in a total assay volume of 60 μl. Following a 72 h incubation at 37 °C, 3% O2, 4% CO2, 30 μl of diluted Sybrgeen reagent was added according to the instructions of the manufacturer (Life Technologies) and fluorescence intensity was quantified using a Biotek Synergy 2 plate reader. Pantothenate competition assays were performed by combining compounds with different concentrations of calcium-d-pantothenate (SigmaAldrich). Activity against asexual blood-stage parasites from a panel of drug resistant strains was determined using 3H hypoxanthine incorporation assays as described before (*29*). Gametocyte viability assays were initiated by inoculating a culture flask with 1% asexual blood-stage parasites in 5% haematocrit in RPMI1640 medium with 10% human serum. From day 4 to day 9 after inoculation, cultures were treated with 50 mM N-acetylglucosamine to eliminate asexual blood-stage parasites. At day 11 post inoculation, gametocytes (predominantly stage IV) were isolated by Percoll density gradient centrifugation as described previously (*30*). Gametocytes were seeded at a density of 5000 cells/well in a 384 well plate and combined with compound diluted in DMSO and subsequently in RPMI1640 medium to reach a final DMSO concentration of 0.1% in a volume of 60 μl RPMI1640 medium with 10% human serum. Following a 72 h incubation at 37 °C, 3% O2, 4% CO2, 30 μl of ONE-Glo reagent (Promega) was added and luminescence was quantified using a Biotek Synergy 2 reader. Effect of compounds on development of *P. falciparum* liver stages in human primary hepatocytes were analyzed essentially as described before (*31*). Briefly, 50.000 human primary hepatocytes were seeded in collagen-coated 96 plates according to the suppliers protocol (Tebu-bio) and combined with compounds serially diluted in DMSO and culture medium to achieve a final DMSO concentration of 0.1%. Hepatocytes were infected with 50,000 *P. falciparum* NF54 sporozoites per well and developing liver schizonts were visualized 4 days after infection by a DAPI nuclear stain and-hsp70 immunostaining as described before (*31*).

For asexual metabolomics studies all parasites were grown in standard RPMI1640 containing ~1-M pantothenic acid and supplemented with 0.25% Albumax II (Gibco) unless otherwise noted. 3D7 was cultured and magnetically purified as previously described (*22*) and NF54 was used for all gametocyte metabolomics studies. Induction (Day −2) of NF54 gametocytes for metabolomics was performed by the addition of 1 volume of minimal fatty acid RPMI (15 μM palmitic and oleic acid only) to a culture of ~10% late ring stage parasites (22-26hpi). This addition was performed without removing the spent media from ~24 hours of growth (1X spent media:1X minimal fatty acid media). On Day-1 the culture was split to ~2% parasitemia with fresh media when the parasites reach the early schizont stage (~38hpi). On the following day (Day +1) a smear was made and the media was replaced to ensure healthy rings are present, a population of which are committed to gametocytogenesis. On Day +2 media was replaced with 2X volumes of media containing 20U/mL Heparin to kill off the replicating asexuals. Heparin media changes (1X volume) were performed on Day +3 and +4. Normal media changes resumed on Day +5 and +6. On Day +7, stage III/IV NF54 gametocytes were purified for metabolomics using the same protocol listed above for 3D7 asexuals.

### Pulse-chase replication assays

Uninfected red blood cells were preloaded with 100μM compound (MMV258, CLQ or DHA), incubated at 37°C and washed thoroughly to wash out the compound. Late trophozoite to schizont stage parasites were magnetically purified and diluted in preloaded uninfected red blood cells to a parasitemia of 0.83%, 3% haematocrit in a total assay volume of 100μl. Following a 72 h incubation at 37 °C, 3% O2, 4% CO2, 100μl of diluted Sybrgeen reagent was added according to the instructions of the manufacturer (Life Technologies) and fluorescence intensity was quantified using a Synergy HTX multi-mode reader (Biotek). Metabolomics pulse-chase assays were performed by seeding uninfected RBCs into a 15mL flask at a density of 1x10^8^ cells/mL in RPMI1640 media (~1% hematocrit). RBCs were pre-incubated for 1 hour before treatment with 1μM MMV689258 (<0.01% DMSO) or no drug for 3 hours. After 3 hours, triplicate 1mL samples were collected from each condition for metabolite extraction (see extraction protocol below). The remaining culture was centrifuged, washed with 1X volume of RPMI, resuspended to the original RBC density, and seeded into a 12-well plate at 1x10^8^ cells/well in 2.5mL of RPMI media (~0.4% hematocrit). Additional triplicate extractions were performed for each condition, as below, at 24 hour intervals post-washout, up to 72 hours post, without subsequent media changes.

### Selection of drug-resistant parasites and fitness assays

*Plasmodium falciparum* strain NF54 was cultured in a semi-automated system as described above and exposed to 30 nM (3 times the IC_50_) CXP18.6-052 to select for drug-resistant parasites. After reappearance and stable growth of surviving parasites, cultures were exposed to 10 times the IC_50_ (90 nM) and subsequently to 50 times the IC_50_ (450 nM). Following stable growth of surviving parasites, limited dilution was performed in 96 wells plates to isolate single cell clones. Drug sensitivity of the resulting clones was analyzed by parasite replication assays as described above. Fitness assays were performed according to Peters et al. (*32*). Briefly, NF54 and a mutant parasites were mixed at a ratio of 1:1 and subcultured biweekly for 4 weeks in the absence of drug pressure. Samples were taken at regular intervals and the relative presence of wildtype and mutant AcCS alleles was assessed by multiplex PCR using primers MWV511 (5′-GGAAGAGATATAGATGGTACTGC-3′) and MWV519 (5′-TCTACCACAATCTGGATATGT-3′) to detect wildtype (T627) AcCS and primers MWV516 (5′-CTACAGTTATTTTTTCTTCTATACCAG-3′) and MWV512 (5′-GATGATTTCCATATACTGTACGTAAC-3′) to detect the mutant (627A) allele. PCR conditions were optimized using a DNA standards comprising wildtype and mutant genomic DNA at preset ratios ranging from 30:1 to 1:30. PCR products were separated on a 1.5% agarose gel and relative quantities were analyzed by densitometry using the FIJI software packag Signals were corrected for background and relative intensities were expressed as a percentage the total amount of amplified DNA.

### CRISPR-Cas9 gene editing

In order to validate the mutations identified by drug-selection, we generated the T627A point mutation in acetyl-CoA synthetase in NF54 parasites using the CRISPR-Cas9 system. Briefly, Cas9 was expressed from the pUF1-Cas9 plasmid (*33*) and the donor template and guideRNA were provided on a pDC2-based *hDHFR* plasmid (a kind gift from Marcus Lee). The donor template was homologous to the target and contained the desired mutation and two additional shield mutations. The guideRNA was expressed under the control of the U6 promoter. Parasites were transfected using an adapted protocol that was previously described (*34*). In short, red blood cells were transfected with both plasmids by electroporation (310V, 950μF) and a synchronised trophozoite culture was added to the DNA-loaded cells. After 1 day, parasites were selected for 5 days using DSM1 (Merck) and cultured until they recovered. Subsequently parasites were selected with 10xIC5_0_ of MMV689258 for 3 days and cloned by limiting dilution. Integration was confirmed by Sanger sequencing.

### Hepatocyte culture and cytotoxicity test

Human primary hepatocytes (Xenotech 098H1500.H15B+) were thawed and plated in a collagen coated 96-well plate in Williams E (Gibco 32551087) supplemented with 1% PenStrep (Gibco 15140-122), 1% Fungizone (Gibco 15290026), 10% hiFBS (Gibco 10270-106), 0,1 IU/ml insulin (Sigma I2643) and 7 μM hydrocortisone hemisuccinate (Sigma H2270) and refreshed daily. After two days of culture compounds were added and incubated in a humidified incubator at 37°C and 5% CO2 for 78 hr (refreshed daily) followed by an additional 18 hr incubation with 700 μM Resazurin (Sigma 199303). Fluorescence was read at Biotek Synergy 2 and data was analyzed in Graphpad Prism version 5.03.

### *In vitro* pharmacokinetics

Plasma protein binding and *in vitro* hepatocyte metabolism data were obtained through a commercial service using their standard protocols (TCG Lifesciences, Kolkata, India). CYP450 inhibition data and CaCo transport data were generated by the Center for Drug Candidate Optimisation at Monash University, Australia using previously described methods (*35*).

### *In vivo* pharmacokinetics

Blood pharmacokinetics in male BALB/c mice were studied by intravenous injection of 3mg/kg compound formulated in 5% (v/v) DMSO and 0.9% (w/v) NaCl, and oral gavage of 30 mg/kg compound formulated in 5% (v/v) DMSO, 0.1%(v/v) Tween 80 and 0.5% (w/v) carboxymethylcellulose in water. Animals were fasted 4 hours pre-and 2 hours after dosing. At timepoints indicated in the figures, approximately 50 μl of blood was collected into heparinized capillary tubes by piercing the saphaneous vein with a needle. Plasma was collected following centrifugation at 1640xg for 5 min at +4°C within half an hour of collection. Plasma samples were stored at-20°C until bioanalysis. Quantitation of compound levels was provided by a commercial service (TCG Lifesciences, Kolkata, India) using their standard LC/MS protocols. Plasma pharmacokinetics in male Sprague-Dawley rats was studied using identical dosing and analyses schemes. Rats were fasted 12 hrs pre-and 2 hrs after dosing.

Biliary and renal clearance was assessed by intravenous dosing of 3 mg/kg compound formulated in 2% (v/v) DMSO, 0.9% (w/v) NaCl to male Han Wistar rats. Bile was collected from bile-duct cannulated rats for 0-1 hr, 1-3 hr, 3-8 hr, 8-24hr and 24-48 hr after dosing. Urine was collected for 0-6 hr, 6-24 hr and 24-48 hr after dosing from rats housed in metabolic cages. Bioanalyses were performed by a commercial supplier (XenoGesis Ltd) using their standard protocols.

### *In vivo* efficacy

Reduction of existing parasitemia *in vivo* was investigated using a humanized mouse model for *P. falciparum* infection. Female NODscidIL2Rγ^null^ mice were engrafted by daily intravenous injection of 0.6 ml of human blood for 11 days. Subsequently, mice were infected with *P. falciparum* strain Pf3D70087/N9 (*36*) by injecting 2.10^7^ parasites in a volume of 0.2 ml. Four days post infection, mice were treated with either vehicle control, MMV689258 or chloroquine. To this end, compounds were formulated in 70% Tween-80 (d= 1.08g/ml) and 30% ethanol (d=0.81g/ml), followed by a 10-fold dilution in H_2_O and administered by oral gavage. Parasitemia was followed by daily collection of 2 μl tail blood. The hematocrit was determined by FACS and parasitemia by microscopy on >10,000 red blood cells as described before (*36*). For pharmacokinetic analyses of samples from the SCID mice, peripheral blood samples (20 μl) were taken at different times as indicated in the figure legends, mixed with 20 μl of ultrapure H2O, immediately frozen on dry ice and stored at −80 °C until analysis. Blood samples were processed under liquid-liquid extraction conditions and analyzed by LC-MS/MS for quantification using a commercial service using their standard protocols (Swiss BioQuant, Basel, Switzerland).

Compound effects on parasite transmission to the mosquito vector were assessed by luminescent Standard Membrane Feeding Assays. To this end, mature stage V gametocyte of transgenic reporter strain NF54-HGL were incubated for 24 hours with compound prior to mosquito feeding as described before (*20*). Similarly, gametocytes from drug-resistant clone F49C11 were fed to *A. stephensi* mosquitoes to assess whether the mutations transmits to the mosquito vector.

### Metabolomics

Extraction and analysis was performed as previously demonstrated (*22*). Briefly, trophozoite stage 3D7 parasites were magnetically purified, allowed to recover in RPMI1640 (0.25% Albumax II) at 0.4% parasitemia (1x10^8^cells/sample), and treated with drug (10X IC_50_) for 2.5 hours. Following treatment parasites were pelleted, washed with mL of ice-cold 1X PBS, and extracted using 1 mL of ice-cold 9:1 MeOH:Water, containing the internal standard ^13^C_4_,^15^N_1_-Aspartate. Supernatants were clarified before drying under nitrogen, followed by resuspension in HPLC grade water containing 1μM chlorpropamide to 1x10^6^parasites/μl for LC-MS analysis. 10 μl was injected on a Thermo Exactive Plus Orbitrap mass spectrometer for LC-MS-based targeted metabolomics (modified from Lu W et al 2010 (*37*). The modifications were made to increase the separation and detection capabilities of late eluting compounds and are as follows: gradients are 0 min = 0%B: 5 min = 20%B: 7.5 min = 55%B: 15 min = 65%B: 17.5 min = 95%B: 21min = 0%B and mass filters are 0-5 min 85-800m/z; 5-6 min 100-800m/z; 6-9.5 min 85-800m/z; 9.5-15.5 min 110-1000m/z; 15.5-22.5 min 250-1000m/z (the last 2.5 minutes are not scanned during re-equilibration). All Orbitrap data was acquired using Phenomenex columns 00D-4387-B0 from batch 5380-0025. Data was processed using Maven as previously described from a targeted metabolite list (*22*) and using a ±10 ppm m/z window and/or a 1 minute RT window from the data provided in Table S2. Post-processing was performed in Microsoft Excel using blank value imputation for compounds at/below background (i.e. avoiding values-0). Graphics were generated using a combination of Excel, R, and Adobe Illustrator. Gametocyte metabolomics was performed as above with a 1 μM treatment concentration, however, the seeding density was 5.5x10^7^ parasites/sample with the same injection volume and concentration. Isotopically labeled standards were purchased from Cambridge Isotope Laboratories, Inc. (Tewksbury, MA).

Saponin lysis was performed as previously demonstrated (*38*) using magnetically purified parasites and parasites were deemed viable through Trypan Blue staining and their ability to metabolize labeled pantothenic acid. Isolated parasites were washed with PBS and resuspended in pantothenate-free RPMI1640 (0.25% Albumax II) (Gibco) to the same densities as above. For analyses of the effects of compounds on CoA metabolism in human primary hepatocytes, cells were cultured in a 12-well plate as described above and 1.4x10^6^ cells were incubated with 1μM compound for 3.5 hours. Cells were then washed with ice-cold 1X PBS, and extracted using 1 mL of ice-cold 75:30 MeOH:Water containing the internal standard ^13^C_4_,^15^N_1_-Aspartate. Additional processing was performed as with parasite samples and the injection concentration was ~2.7x10^4^cells/μl. Primary hepatocytes were extracted in technical duplicate and pooled for a single LC-MS injection due to sample limitations.

### Whole genome sequencing

Genomic DNA was isolated from parasites a DNeasy Blood & Tissue Kit according to the instructions of the manufacturer (Qiagen). Library preparation was performed as previously demonstrated (*39*). Briefly, Illumina barcoded DNA quality was assessed using an Agilent 2100 Bioanalyzer and Qubit before samples were sequenced using an Illumina HiSeq 2500 system (Penn State Genomics Facility). Sequencing outputs were uploaded into a local instance of Galaxy and quality assessed by FASTQ. Sequences were trimmed using trimmomatic and mapping was performed using the 3D7 reference genome (www.plasmodb.org) and NGS Mapping (BWA-MEM) program. Files were filtered and converted to SAM/BAM for variant analysis by FreeBayes. Variants were identified and annotated using the SNPeff tool followed by further selection using the Select variants and variant filtration tools. Visualization of the alignment and unique reads was performed using the Integrative Genomics Viewer. Mutations in the coding regions which were unique to the resistant line(s) but not the parental NF54 were denoted as significant contributors. Read depth was manually assessed for genes of interest (pantothenate/CoA biosynthesis pathway) to ensure there was no copy number variation between the tested parasite lines.

### Generation of recombinant PfPANK1

Recombinant PfPANK1 used for immunization was obtained by cloning the cDNA corresponding to ORF2 that encodes the full-length PfPanK1 protein into the baculovirus transfer vector pFastBacHT in frame with an N-terminal 6-histidine purification tag. PfPanK1 protein was produced with the baculovirus expression system at the St. Jude Childrens Research Hospital Protein Production Facility. Bacmid production, transfection, and baculovirus amplification were carried out according to the manual (Invitrogen). Suspension cultures of *Sƒ*9 cells were cultured in SFX-insect serum-free media to a density of 2 x 10^6^ cells/ml and infected at a multiplicity of infection of 10. Cultures were gently shaken for 3 days at 28°C, harvested and resuspended in a buffer containing 50 mM Tris, pH 7.9, 500 mM NaCl, and 10% glycerol and lysed by microfluidization. Cell lysates were clarified by centrifugation at 20,000 rpm and soluble PfPanK1 was purified from the supernatant by Hitrap Ni^2+^metal affinity chromatography and eluted with 50 mM Tris, pH 7.9, 500 mM NaCl, 10% glycerol and 1 M imidazole. Purified PfPanK1 was dialyzed into 50 mM Tris, pH 7.9, 500 mM NaCl and 10% glycerol flash frozen in liquid nitrogen, and stored at −80 °C until further use.

### Generation of recombinant human PANKs

Recombinant human PANK1, PANK2, and PANK3 were cloned and expressed using a bacterial expression system. The gene encoding human PANK1-was mutagenized using the QuickChange site-directed mutagenesis kit (Stratagene) with primers that added a NheI restriction site to the NH2-terminus and a XhoI restriction site to the COOH-terminus. The PCR product was cloned into pET28a to obtain NH2-terminal His6 tag fusion protein and the resulting plasmid was transformed into the *E. coli* BL21(DE3) (Stratagene) expression strain. The cells were grown to mid-log phase and the protein was induced with 1.0 mm IPTG for 18 h at 16 °C. The PANK1-protein were purified using nickel-NTA affinity column and eluted with 50 mM Tris, pH 7.9, 500 mM NaCl, 10% glycerol, 300 mM Imidiazole. The purified protein was dialyzed overnight at 4 °C in 20 mM Tris, pH7.5, 300 mM NaCl and stored at −80 °C in equal volume glycerol until further use. The genes encoding the mature form of human PANK2 (residues 141-570) and the catalytic core domain of human PanK3 (residues 12-368) were similarly cloned and purified as previously reported (*40*, *41*).

### Generation of a polyclonal anti-PfPANK1 serum and immunoprecipitation of PfPANK1

Rabbits were immunized with recombinant PfPANK1 according to standard procedures of the manufacturer (Eurogentec, Seraing, Belgium). Immunoglobulins were absorbed on protein A/G sepharose and used to isolate PfPANK1 from parasite lysates. PANK assays, as described below, were performed on the immunoprecipitated material and the depleted parasite lysates.

### Parasite lysates for PfPANK assays

Aynschronous blood-stage parasites from *P. falciparum* strain NF54 were released from the red blood cells by lysis with 0.06% saponin in PBS for 5 minutes on ice. Parasites were pelleted by centrifugation (10 minutes at 4000xg), washed with PBS and lysed in 50 mM NaF, 20 mM Tris-HCl (pH 7.5), 0.1% Triton-X, 2 mM dithiothreitol, 2 mM EDTA and 1% (v/v) Halt Protease Inhibitor Cocktail (Thermo-Fischer Scientific, Waltham, MA, USA). Suspensions were then sonicated 6 times 3 seconds at an amplitude of 16 microns peak-to-peak. Sonicated samples were centrifuged at 15000 rpm for 5 minutes at 4°C. The supernatant was then used in the activity assay.

### PANK assay

PANK activity was monitored using a radioactive labeled pantothenate kinase assay. The reaction mixtures contained 10 mM MgCl_2_, 2.5 mM ATP, 5 *μM* 14C-labeled pantothenate, 100 mM Tris-HCl, pH7.4, the to be tested pantothenamide and enzyme (freshly prepared Plasmodium falciparum lysate, rec. protein in a total volume of 40 μl. Reactions were initiated after addition of enzyme and incubated at 37 degrees for 60 minutes. Reaction were terminated with 4 μl of a 10% acetic acid solution in 95% ethanol. Samples were loaded on DEAE filter discs (GE Healtcare) and washed thoroughly in 1% acetic acid solution in 95% ethanol. Phosphorylated pantothenate will stay on the filter, while the unphosphorylated pantothenate will be washed away. After the discs were dried, they were transferred into scintillation vials containing 3 ml ScintiSafe 30% Cocktail (Fischer Scientific, Hampton, NH, USA). Radioactivity in each vial was counted using a Tri-Carb 2900TR Liquid Scintillation Analyzer (Packard Bioscience, Boston, MA).

## Acknowledgements

This work was supported by National Institutes of Health Grants GM062896 (SJ), Cancer Center Support Grant CA21765 (SJ), Medicines for Malaria Venture grant RD/14/0019 (KJD, ML, JS), Burroughs Wellcome Fund Investigators in Pathogenesis of Infectious Disease (PATH) Award (ML), NIH Ruth Kirschstein National Research Service Award (NRSA) Individual Postdoctoral Fellowship (F32) AI124507 (EA) and the American Lebanese Syrian Associated Charities (SJ). TWAK is supported by the Netherlands Organization for Scientific Research (NWO-VIDI 864.13.009). The authors wish to thank Jeremy Burrows for advice and discussion and the Lygature product development partnership for help with project management. A cDNA encoding PfPANK1 was provided to S.J. by C. Ben Mamoun, Yale University. Marcus Lee is kindly acknowledged for contributing a pDC2-based gene targeting vector. The authors would also like to thank the Huck Institutes of Life Sciences core facilities-Penn State Penn State Genomics Core Facility and Penn State Metabolomics Core Facility-University Park, PA. The Illumina HiSeq 2500 (Penn State) was purchased with an NSF-MRI award DBI-1229046. In particular the authors thank Drs. Andrew Patterson and Phil Smith from the Penn State Metabolomics Core for technical and metabolomics expertise and Susan Charman and Karen White from the Center for Drug Candidate Optimisation of Monash University for in vitro DMPK assays.

